# AI-Driven System for Large-Scale Automated Collection of Mouse Profile Images

**DOI:** 10.1101/2025.03.27.645835

**Authors:** Naomitsu Ozawa, Yusuke Sakai, Yusuke Sakai, Chihiro Koshimoto, Seiji Shiozawa

**Affiliations:** Institute for Disease Modeling, Kurume University School of Medicine, 67 Asahimachi, Kurume City, Fukuoka 830-0011, Japan; Division of Bio-resources, Department of Biotechnology, Frontier Science Research Center, University of Miyazaki, Kihara 5200, Kiyotake, Miyazaki 889-1692, Japan; Laboratory Animal Center, Keio University School of Medicine, 35 Shinanomachi, Shinjuku-ku, Tokyo 160-8582, Japan

**Keywords:** Artificial Intelligence, Facial Expression Recognition, Image Analysis

## Abstract

As with human communication, recent studies have revealed that animals convey a substantial amount of information through their facial expressions. In these studies, artificial intelligence (AI) technologies have been increasingly employed to analyze animal facial image data. However, collecting large amounts of facial image data for such studies has been labor-intensive. In this study, we developed a system that automatically recognizes and saves the faces of freely moving mice using AI-based object detection and image classification technologies. Through validation experiments, we confirmed that the system can detect, classify, and save a variety of mouse profiles with high accuracy. To further expand the versatility of the system for diverse research applications, the technology has been improved to include a feature for determining mouse sex based on their profiles, leveraging AI algorithms for this purpose. A small dataset was used to evaluate the performance of the sex determination system, yielding 100% accuracy for both male and female classifications. This application enables researchers to efficiently collect facial image data, providing high-quality datasets suitable for AI training. Consequently, the efficiency of facial expression analysis in mice is significantly improved. Importantly, this technology is not limited to mice and has the potential to be applied to other animal species and a wide range of research fields, offering promising potential for diverse applications.

## Introduction

Facial expressions are a pivotal form of communication in humans, expressing thoughts and emotions [1]. In recent years, multiple studies have revealed that animal facial expressions also convey substantial information, and the analysis of these expressions has seen significant advancements [2]. The Animal Facial Action Coding System (AnimalFACS), a tool designed to analyze animal facial expressions, was developed to systematically analyze animal expressions by examining the movements of facial muscles [3]. For instance, the DogFACS is a standardized system for analyzing and categorizing facial movements in canines based on their underlying muscular activity [4,5]. It assists researchers and trainers in objectively assessing canine expressions related to emotions and communication. Similarly, CalliFACS has enabled comprehensive documentation and analysis of facial movements in common marmosets, a nonhuman primate species increasingly recognized as a valuable model in experimental research [6], making it an essential tool for studying social behavior and emotional expression [7].

Furthermore, it has been demonstrated that the degree of pain can be estimated from the facial expressions of mice, which represents a significant discovery contributing to the enhancement of animal welfare [8]. Specifically, the Mouse Grimace Scale (MGS), based on subtle facial expression changes in mice, is extensively utilized for pain assessment. A similar approach has been applied to rats [9], rabbits [10], and cats [11], where grimace scales have been developed to assess facial expression changes in these species. These scales have been incorporated into objective methods for pain assessment [12,13].

On the other hand, accurate assessment requires expertise, and since it is a subjective measure, there is considerable variability in the results. As a result, there is an increasing demand for automation to ensure consistency and objectivity in assessments [12]. Furthermore, it has become evident that mice not only convey pain but also contain more advanced information in their facial expressions [14]. Research has been conducted to determine whether mice display specific facial expressions in response to emotionally significant stimuli, and whether these expressions reflect fundamental characteristics such as emotional intensity, value, flexibility, and persistence [15,16]. In addition, there have been reports of a novel method that combines mouse facial expression analysis with two-photon calcium imaging, which has enabled the identification of single neurons in the insular cortex— a brain region believed to be involved in human emotional experiences [17]. These neurons were found to be closely correlated with specific facial expressions. This finding suggests that mouse facial expressions may carry more information than previously assumed, and further analysis is expected. These studies suggest that facial expressions are not merely reflections of brain activity; rather, they play a pivotal role in enhancing our understanding of behavior and social interactions.

However, conducting such analyses of animal facial images requires the manual extraction of face images from videos of animals exhibiting free-ranging behavior to serve as the original data [14]. This approach demands a large volume of data, which has proven to be a significant challenge for researchers [2]. Furthermore, the manual extraction process can introduce inconsistencies and issues with reproducibility, potentially undermining the reliability of research findings. As a result, there is a pressing need to implement more efficient and standardized methodologies to overcome these challenges.

Recent research has employed a method for acquiring facial images by head-fixing mice for deep learning analysis [17–19]. Studies have reported that immobilization using a head-fixation apparatus may impact the natural behavior and facial expressions of mice, requiring an acclimation period of 10 to 25 days to minimize its effects [20]. During this period, mice undergo specialized training to acclimate to the fixation apparatus, a process that demands significant time and effort from researchers, thereby reducing overall research efficiency.

To address these issues, the objective of this study was to develop an application that automatically detects, classifies, and stores the faces of mice exhibiting free-ranging behavior in video clips. This approach will not only significantly reduce the burden on researchers and improve data collection efficiency, but also contribute to research efficiency by eliminating the need for acclimation and training associated with fixation. From an animal welfare perspective, this method is more favorable, as it minimizes stress on the animals. Furthermore, the integration of an artificial intelligence (AI) system to evaluate stored profiles is expected to expand the application’s functionality to a range of tasks. For instance, it could be used for developing new analysis methods based on facial morphological features. In this study, we demonstrated the feasibility of applying this function to determine mouse sex, a task that is often difficult for humans. This application aims to automate the evaluation of pain based on the Grimace Scale, as well as the analysis of various facial expressions associated with specific emotional states.

## Materials and Methods

### Ethical statement

All animal experiments were performed in accordance with the guidelines for laboratory animals set forth by the National Institutes of Health and the Ministry of Education, Culture, Sports, Science and Technology (MEXT) of Japan and were approved by the Kurume University Animal Experiment Committee (approval number: 2024-127) and the University of Miyazaki Animal Experiment Committee (approval number: 2023-527).

### Animals

C57BL/6J mice were purchased from CLEA Japan, Inc. (Tokyo, Japan). C57BL/6N, C3H/He, and ICR mice were purchased from Japan SLC, Inc. (Shizuoka, Japan). Video recordings of *Apodemus speciosus* were made using animals bred and maintained at the University of Miyazaki.

### Video Recording

Three cameras were used: an iPhone 7 (Apple Japan G.K., Tokyo), a HOZAN L-834 near-infrared camera (Hozan Co., Ltd., Osaka), and a SONY ZV-1 II (Sony Group Corporation, Tokyo).

The video recordings were captured in Full HD resolution using the iPhone 7 and HOZAN L-834 near-infrared camera, and in 4K UHD resolution with the SONY ZV-1 II. The frame rate for all recordings was consistently set to 30 fps.

The recordings were conducted inside a custom-made recording box (150 mm × 100 mm × 100 mm). For the development of the head detection AI and profile classification AI, video recordings of four mouse strains (C57BL/6N, C57BL/6J, C3H/He, and ICR) were made at Kurume University. Ten males and ten females of C57BL/6N, and five males of C3H/He were recorded using the iPhone 7. Five females of ICR were recorded using the SONY ZV-1 II. Additionally, one male of C57BL/6J was recorded using the near-infrared camera. Some videos of ICR and *Apodemus speciosus* were recorded at Miyazaki University using the SONY ZV-1 II.

For the development of the sex determination AI, 10 males and 10 females of C57BL/6N were recorded using the iPhone 7. Three 1-minute video recordings were obtained for each animal.

### Development Environment

Python was used as the primary programming language. Video and image processing were primarily implemented using OpenCV. For deep learning, PyTorch was used as the primary framework for object detection AI, while TensorFlow and Keras were employed as the primary frameworks for classification AI. Deep learning tasks were mainly performed on a workstation. The main specifications of the workstation included Ubuntu 22.04 LTS as the operating system, an Intel Core i7 13700K CPU, an NVIDIA RTX 4070 Ti GPU, and 128 GB of RAM (hereafter referred to as RTX-WorkStation). For inference, a standard-performance PC, the 13-inch MacBook Pro (2020 Apple Silicon M1 model, hereafter referred to as M1MacBookPro), and the high-performance RTX-WorkStation used for training were employed.

### Dataset preparation

Mouse head annotations were performed using Microsoft’s VOTT2 tool. A total of 710 images of C57BL/6N, 978 near-infrared images of C57BL/6J, 1,594 images of ICR, and 486 images of *Apodemus speciosus* were annotated. Augmentation was applied by randomly adjusting brightness, exposure, and noise within the bounding boxes, leading to a three-fold increase in dataset size.

The developed head detection AI was then used to extract head images from each video. These extracted head images were categorized and labeled as either side profile (“OK”) or non-side profile (“NG”). Image augmentation was applied to this dataset (Table S1), resulting in 5,889 images for accuracy evaluation, 117,398 images for training, and 29,082 images for validation (Table S2).

For the sex determination AI, side-profile images were extracted from videos of the animals using the developed application. Data cleaning was performed to remove images with significant issues, such as missing large portions of the face. Image augmentation was applied to this dataset (Table S1), resulting in 1,044 images for accuracy evaluation, 110,182 images for training, and 27,450 images for validation (Table S3).

### Development of head detection AI

To detect the heads of free-moving mice, one of the object detection AIs, Ultralytics YOLOv8 [21], was used. This state-of-the-art (SOTA) model allows for easy training of custom models. First, frames were extracted from the videos, and K-means clustering was applied to obtain diverse frame patterns. A dataset annotated with mouse head locations in these frames was used for training. The model’s performance was evaluated using the mAP50 and mAP50-95 metrics.

### Development of Side-Profile Classification AI and Sex Determination AI

The training scripts were developed using TensorFlow and Keras as the primary frameworks. An additional image augmentation pipeline was implemented in the data input pipeline, allowing various augmentations (see Table S4) to be applied during training. In the final stage of image augmentation, the background was replaced with either a random gray value or random noise with a 50% probability. This approach was implemented based on previous reports indicating that it can improve classification accuracy [22]. Training was efficiently conducted by adding a classifier to the headless MobileNetV3Large [23] model, which is known for being lightweight and high-performance, and then fine-tuning it. The model was trained using “Categorical Focal Cross-Entropy” [24] as the loss function to address class imbalance issues, with the “Adam” optimizer employed. During training, the weights up to the bottleneck blocks preceding Conv_1 were frozen, while the layers after Conv_1 were retrained and fine-tuned (see Table S5).

The model’s performance was evaluated by inferring the accuracy evaluation dataset and generating a confusion matrix of true positives, true negatives, false positives, and false negatives. Performance metrics, including accuracy, precision, recall, and the F1 score (the harmonic mean of precision and recall), were used for evaluation.

For the sex determination AI, validation was also conducted using videos. Inference was performed on videos that were not used during training to determine sex. For sex determination in videos, all detected side-profile images within a single video were evaluated. If classified as male, 1 point was added to the male score; if classified as female, 1 point was added to the female score. In cases where the classification results for males and females were equal, 1 point was added to both scores. After evaluating all detected side profiles, the male and female scores were compared, and the higher score was taken as the sex determination result for the video. As with the model evaluation, a confusion matrix was generated and used for assessment.

### Development of an Algorithm for Determining the Focus of Selected Face Images

To detect image blur using OpenCV, one common method involves calculating the variance of image edges using an edge detection algorithm and judging the image based on that value [25]. In this study, side-profile images were collected, and the Sobel filter (XY), an edge detection algorithm, was applied to calculate the variance values for each image. The distributions of in-focus and out-of-focus images were then compared, and an initial threshold value for focus determination was established.

### A Method for Clustering Collected Face Images

The K-means algorithm was employed to cluster the large number of images obtained through the head detection AI and the side-profile classification AI.

First, to extract feature vectors from the acquired images, a headless MobileNetV3Small model pre-trained on ImageNet was used. This model produces a 7×7×576 output, which was flattened into a 1-dimensional vector with 28,224 elements, thus obtaining the feature vectors.

The extracted feature vectors were then reduced to three dimensions using t-SNE from Scikit-Learn, followed by K-means clustering. After clustering, to avoid obtaining similar patterns of images, the centroid of each cluster was calculated, and the image closest to the centroid was selected from each cluster.

### Background Removal

During clustering, the frame numbers within the video were linked to each frame. The full images corresponding to the frame numbers extracted through clustering were then processed using remBG [26,27], a Python library, to remove the background. After background removal, the side-profile regions were cropped from the processed images to produce the final output.

### Application Development

The head detection AI, focus check function, side-profile classification AI, and clustering function were integrated into a single application. Optimized models for the head detection AI and side-profile classification AI were deployed for each inference platform. When performing inference on Apple Silicon M1 or later, models converted into CoreML format using Core ML Tools were used. For inference on Windows or Linux systems equipped with NVIDIA discrete GPUs, models saved in Keras format were utilized.

The video input and pipeline construction were handled using OpenCV. A pipeline was created to process each video frame sequentially through the head detection AI, focus check function, and side-profile classification AI. Images identified as side profiles within the video, along with their frame numbers, were stored, and the corresponding full-frame images were temporarily saved to storage. These side-profile images and frame numbers were then passed to the clustering process, where dimensionality reduction was performed using t-SNE, followed by K-means clustering.

After clustering, the frame numbers closest to the centroids of each cluster were identified. The temporarily saved full-frame images corresponding to the identified frame numbers were processed using remBG to remove the background. Finally, the background-removed images were passed to the head detection AI to crop the face regions, which were then saved.

As an additional feature, by modifying the pipeline after the side-profile classification AI, the application can be adapted for various other purposes.

### Performance Evaluation

For the performance evaluation of the application, the processing time for each frame was measured, and the average processing time per frame was calculated using a 5% trimmed mean to reduce the influence of outliers.

### Grad-CAM++ Analysis for CNN-Based Sex Determination

Grad-CAM++ was applied to visualize the regions contributing to the CNN’s sex determination. The analysis was performed using the tf-keras-vis library, with Grad-CAM++ applied to the Conv_1 layer of MobileNetV3Large. Heatmaps were overlaid on the input images to highlight the key features used for classification.

## Results

In this study, we first aimed to collect image data of the mouse head under free-moving behavior, where the head is not fixed. To achieve this, we developed an object detection AI to detect the mouse head. The trained model was evaluated using mAP50 and mAP50-95, which are common evaluation metrics for object detection tasks (Table 1). mAP (mean average precision) is a metric used to evaluate the accuracy of object detection predictions. mAP50 represents the average accuracy when the IoU (Intersection over Union) threshold is set at 0.50, while mAP50-95 reflects the average accuracy across a range of IoU thresholds from 0.50 to 0.95. IoU is a measure of the overlap between the predicted and actual regions. For example, if the IoU threshold is set to 0.50, a prediction is considered correct if the predicted region overlaps with the actual region by at least 50%, while an IoU threshold of 0.95 requires a 95% overlap to be considered correct. A mAP50 value of 0.995 indicates that 99.5% of the detected mouse heads were accurately identified. Additionally, a mAP50-95 value of 0.892 means that even under stricter conditions (with a higher IoU threshold), mouse heads were detected with 89.2% accuracy. When inferring from the images of the animal species used for training, the model successfully detected heads with high precision in those species (Fig. 1A-D). A demonstration of the detection process is shown in Supplementary Movie. Furthermore, even in the unlearned C3H/He species, detection was highly accurate (Fig. 1E). These results demonstrate that the object detection AI successfully detected the heads of free-moving mice. Subsequently, an attempt was made to automatically filter out in-focus images among the detected head images. Out-of-focus images often appear in video clips, and there is concern that these low-quality images may negatively affect the accuracy of the analysis. To address this issue, OpenCV’s edge detection function was applied to select only in-focus images, which were then passed to the profile detection AI for further processing.

**Figure 1.**
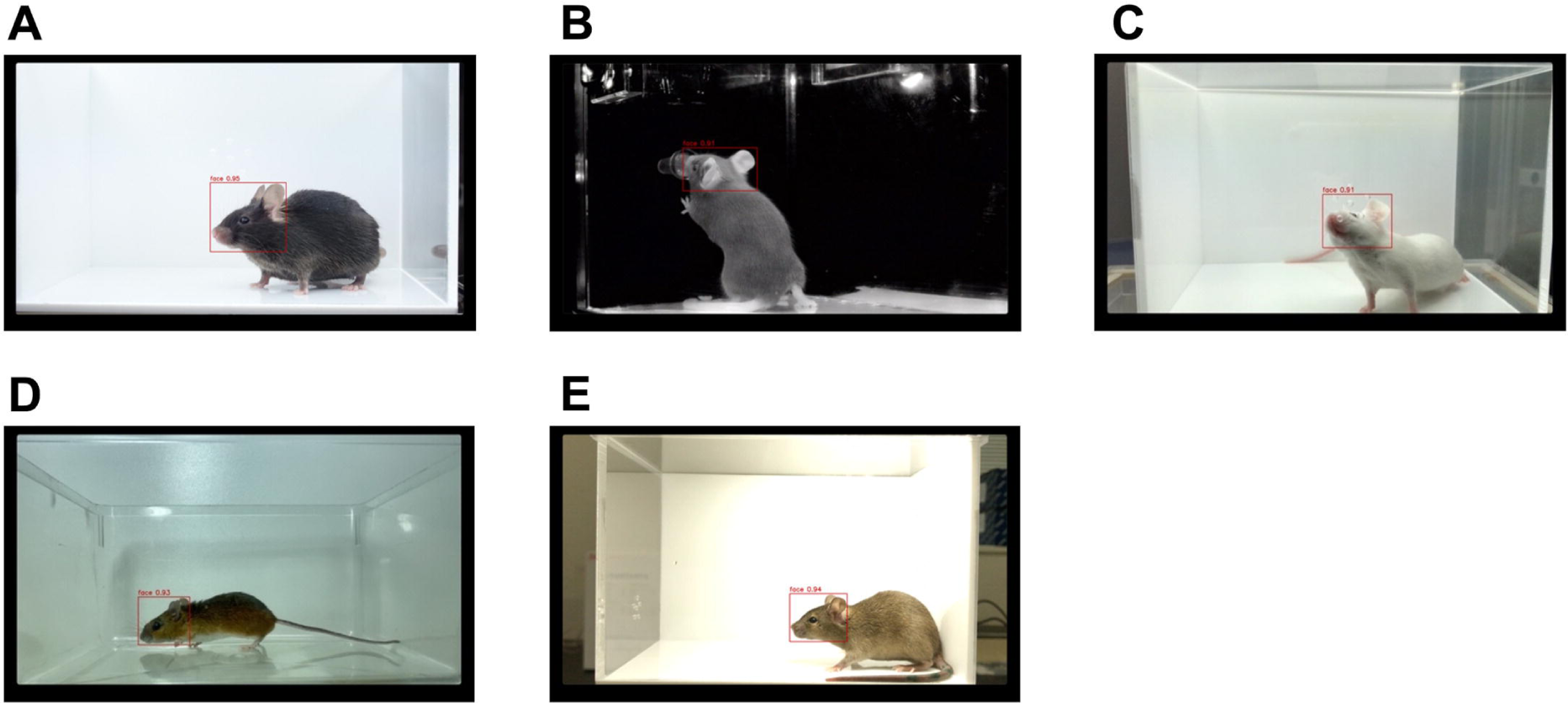
Head detection AI. Head detection performance was evaluated using bounding box (BBOX) annotations on three mouse strains and *Apodemus speciosus* images. (A) C57BL/6J. (B) Near-infrared (NIR) image of a C57BL/6J. (C) ICR. (D) *Apodemus speciosus*. (E) C3H/He. BBOXes indicate AI-detected head regions.

Initially, the dispersion values of profile images extracted from 4K-resolution videos of ICR and C57BL/6J mice were summarized in the figure (Fig. 2A). The distribution of dispersion values for ICR and B6, two mouse strains with different coat colors, suggested that focus determination could be performed around a dispersion value of approximately 2,600. Therefore, the default threshold for focus checking was set to 2,600. However, in the case of B6, which has black fur, we investigated the effect of video brightness. When the exposure value (EV) of the reference video was set to 0EV, it was observed that under brighter conditions (equivalent to +1EV), the outline of the black fur became more accentuated, leading to a higher dispersion value. Conversely, in darker conditions (equivalent to -1EV), the fur outline became less distinct, resulting in a lower dispersion value (Fig. 2B). This finding suggests that the threshold needs to be adjusted based on both the animal’s coat color and the exposure settings during filming. Based on this insight, we have made it possible for users to specify the appropriate threshold value by setting this parameter as an argument in the application specifications.

**Figure 2.**
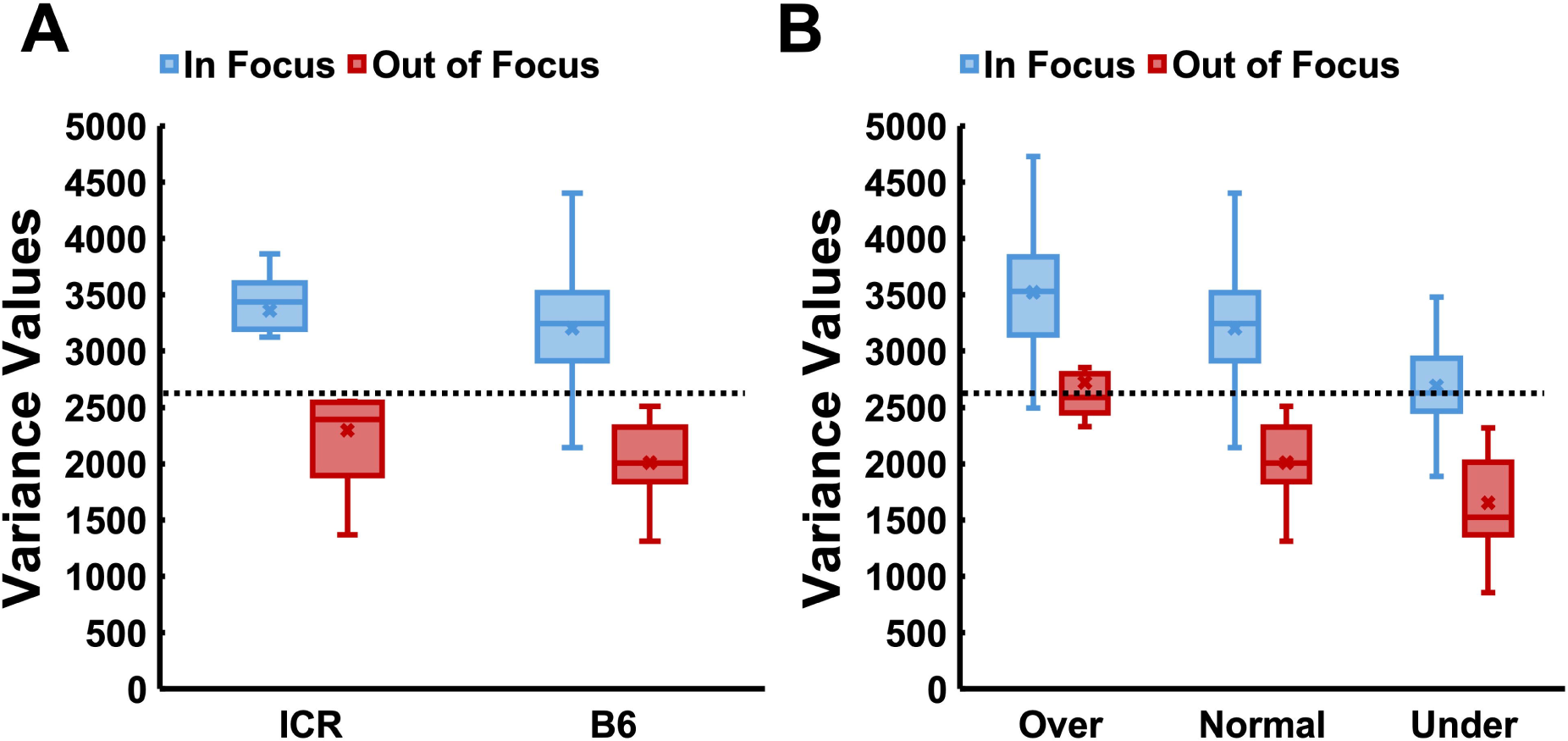
Evaluation of the focus-checking function based on variance values of side-profile images. (A) Variance values of side-profile images for ICR and C57BL/6J (B6) mice with different coat colors. A dotted line indicates the threshold for distinguishing in-focus and out-of-focus images. (B) Variance values of side-profile images for B6 mice under varying exposure levels. A dotted line represents the threshold for distinguishing in-focus and out-of-focus images under normal exposure conditions.

Subsequently, an attempt was made to develop an image classification AI system capable of selecting only the profile images from the obtained mouse head images. To achieve this, the AI was trained as a binary classifier to distinguish between profile and non-profile images. The training was prematurely stopped at epoch 149, with the weights from epoch 129 saved as the optimal model. The accuracy was 0.9495 for the training data and 0.9747 for the evaluation data, while the loss was 0.0082 for the training data and 0.0046 for the evaluation data, demonstrating a high level of accuracy, as shown in Figures 3A and B. After training, accuracy data was predicted for the evaluation phase, and a confusion matrix was calculated, with “OK” designated as positive and “NG” as negative. The matrix showed 1,402 true negatives, 392 false negatives, 98 false positives, and 3,997 true positives (Table 2). Based on this confusion matrix, various evaluation metrics were calculated and assessed (Table 3). First, the “Accuracy” was 91.68%, indicating that the model’s predictions were mostly correct. The “Precision” was 93.47%, meaning that 93.47% of the predicted profiles were indeed profiles. The “Recall” was 78.15%, showing that 78.15% of actual profiles were correctly identified. Finally, the F1-Score, which represents the harmonic mean of precision and recall, was 85.12%, suggesting a good balance between precision and recall. These evaluations collectively demonstrate that the model performs well, particularly in accurately selecting only the profile images.

**Figure 3.**
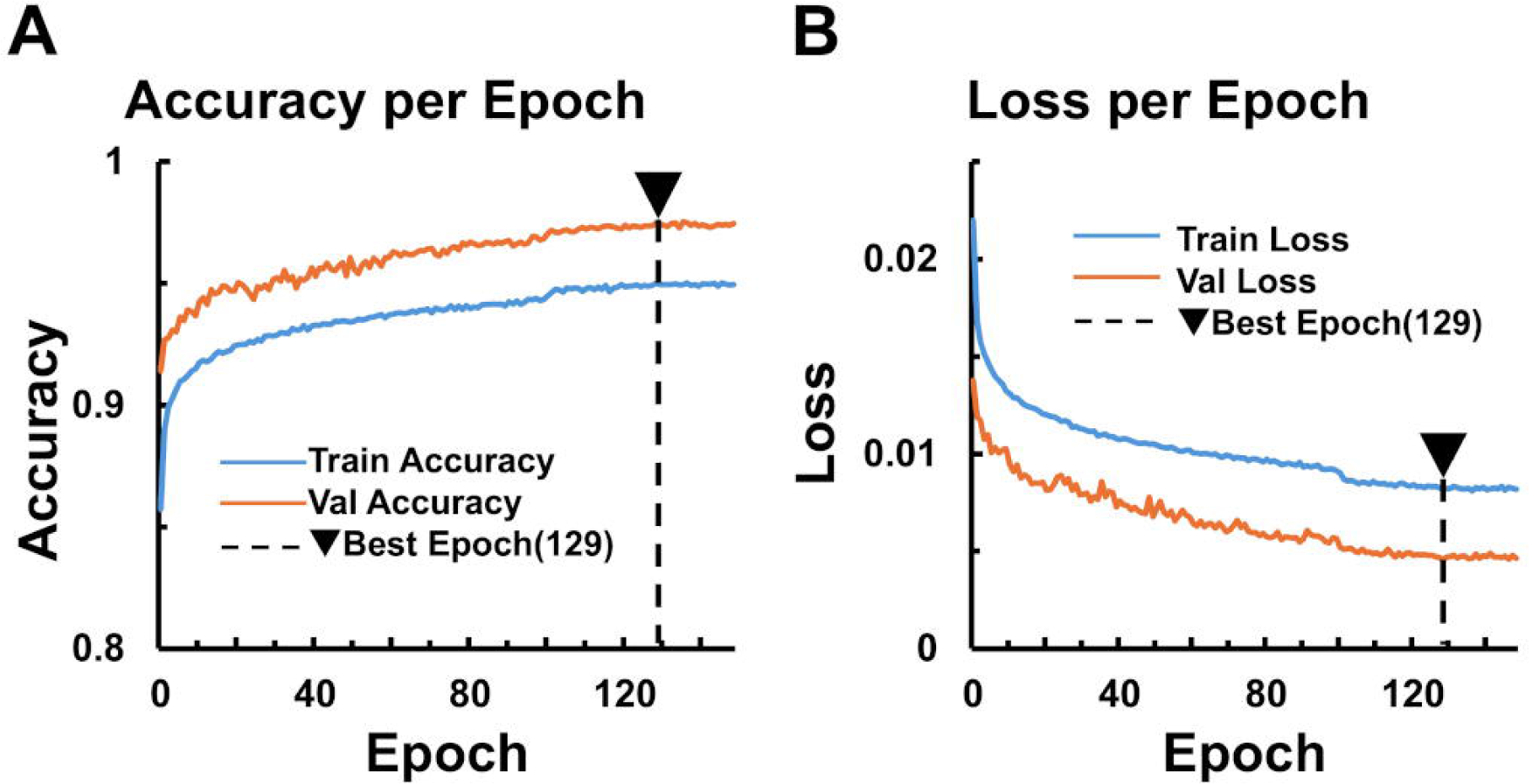
Side-Profile Classification AI. (A) Accuracy per epoch during training and evaluation. (B) Loss per epoch during training and evaluation. Early stopping was triggered at epoch 149, with the model weights from epoch 129 saved as the best. The results were as follows: for the training data, loss = 0.0082 and accuracy = 0.9495; for the evaluation data, loss = 0.0046 and accuracy = 0.9747.

The resulting profile images are extracted from consecutive frames of video, leading to a substantial number of images with high local similarity and a lack of diversity. This was anticipated to negatively impact the training of the image classification AI by introducing bias into the dataset’s characteristics. Therefore, an effort was made to select images with diverse patterns from the collected set by excluding nearly identical images from successive frames.

Initially, attention was given to the feature extraction capabilities of CNNs as a method for obtaining features from the collected images. The small and lightweight MobileNetV3Small headless model, pre-trained on the large ImageNet dataset, is widely used as a backbone in various AI applications. The 7×7×576 output from this headless model was converted into a one-dimensional vector of 28,224 elements, then reduced to three dimensions using t-SNE, followed by clustering using the K-means algorithm.

To obtain representative data for each cluster, a Euclidean distance-based approach was used to select the image closest to the center of each cluster. This method ensured that data remained as unbiased as possible while preserving distance independence from other clusters. To visualize the results of the clustering, the outcomes for K=3, 8, and 16 were displayed in 3D graphs (Fig. 4A), along with the profile image closest to the center of each cluster (Fig. 4B). Furthermore, since a large number of similar images appeared to be clustered in Group C in Fig. 4B, we also extracted the full-body images and frame numbers for Group C (Fig. 4C). These steps allowed us to successfully exclude nearly identical images from consecutive frames, collecting only those with relatively high diversity.

**Figure 4.**
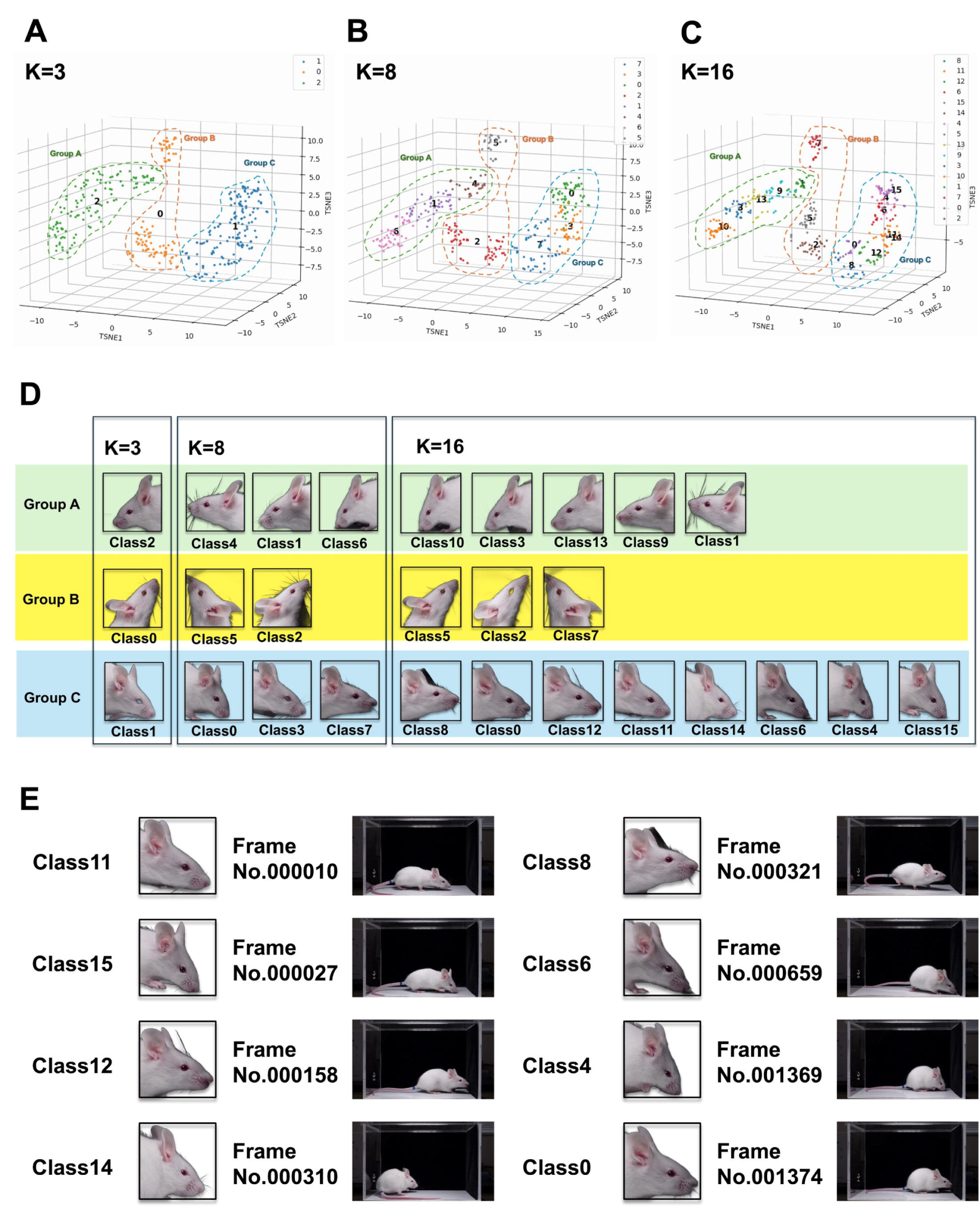
Clustering of side-profile images using K-means and t-SNE. (A) Scatter plots of side-profile images clustered with K = 3 (Left), K = 8 (Middle), and K = 16 (Right) after dimensionality reduction with t-SNE. (B) Side-profile images nearest to the cluster centers for each K value. (C) Full-body images corresponding to the side-profile images in Group C clusters.

Finally, an evaluation was conducted to assess the processing speed of these pipelines and the effect of parallelization. Since the pipelines process one frame at a time, the processing time per frame (Fig. 5A) and the average frame time for a 1-minute video (Fig. 5B) were measured. Measurements were taken using two different execution environments: a standard performance environment (M1 MacBook Pro) and a high-performance environment (RTX WorkStation). The software operates in two modes: one where processing occurs while previewing the bounding boxes and profile AI decision results on the input video, and another where processing occurs without displaying these results. The mean frame time for the prediction pipeline in each environment was calculated using a trimmed mean (5%) to exclude outliers. This approach was necessary because sporadic increases in processing time, such as stuttering at the beginning or middle of the video, sometimes occurred. These instances were considered outliers and were therefore excluded from the data set.

**Figure 5.**
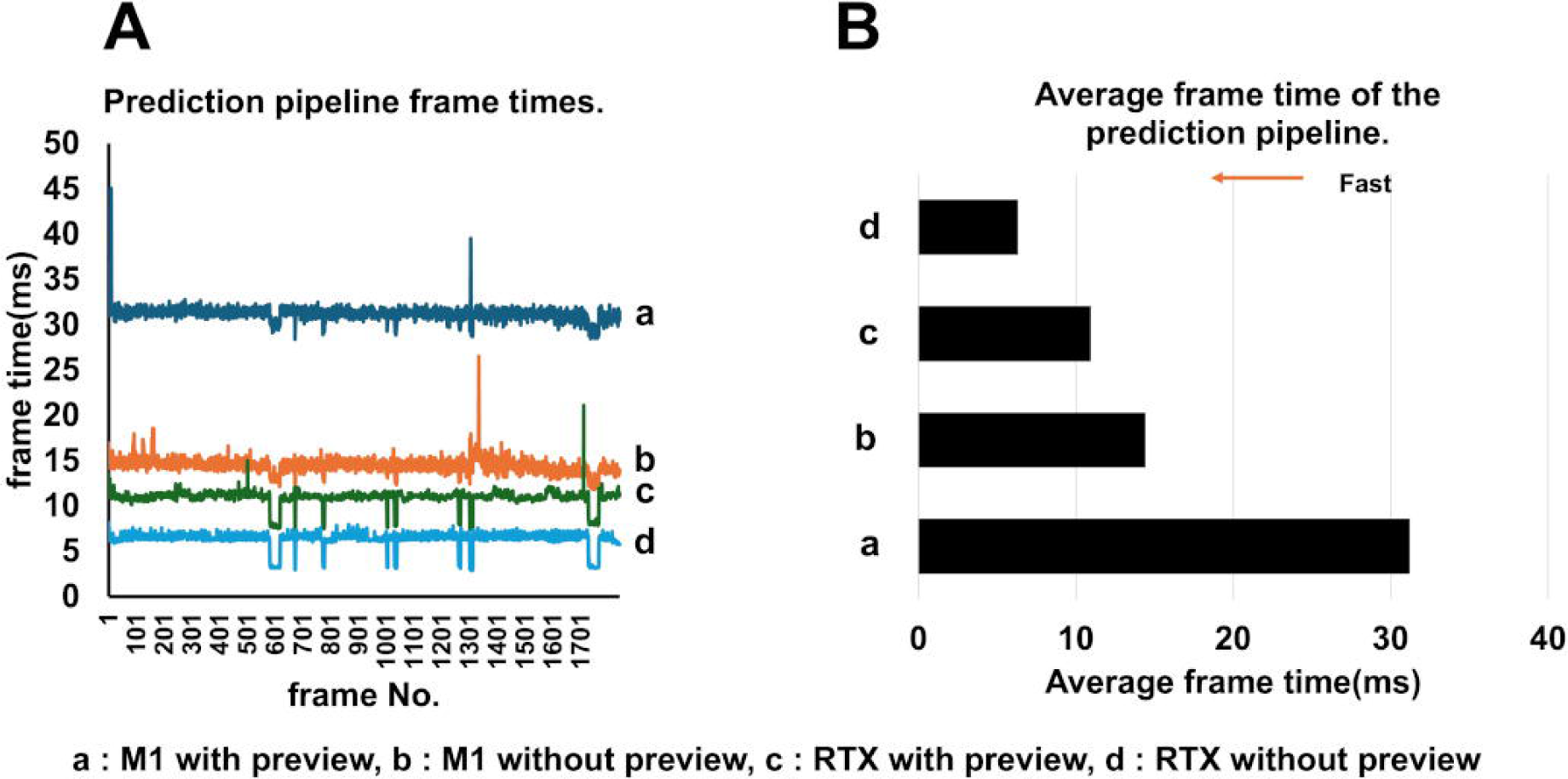
Performance evaluation of the prediction pipeline. (A) Processing time per frame in the prediction pipeline (ms). (B) Average processing time per frame.

Initially, the RTX-WorkStation without preview demonstrated the fastest processing speed, with an average time of 6.3 milliseconds (msec). The RTX-WorkStation with preview followed with an average of 10.9 msec, while the M1 MacBook Pro recorded an average of 14.4 msec without preview and 31.2 msec with preview, indicating sufficient processing power for the task.

Based on the above results, we have successfully developed an application that efficiently collects mouse profile images in this study. This application can be adapted for various research purposes by incorporating an AI trained to judge profiles for different tasks within its prediction pipeline. To demonstrate its versatility, we first decided to investigate whether it is possible to determine the sex of a mouse based on its facial image.

In general, sex determination in mice is typically based on anatomical differences in external genitalia, mammary gland development, and other systemic characteristics, making it challenging to determine sex from facial images alone. However, we believe that AI has the potential to distinguish subtle differences that are difficult to detect visually, making it an ideal tool for demonstrating the value of AI-based facial image analysis. To explore this, we first used this application to collect a small dataset from C57BL/6N images and developed a prototype AI for sex determination. Training employed early stopping, where the weights from the best-performing epoch within the last 20 epochs were saved if the loss value did not improve or increased significantly in the evaluation data. The training was terminated at epoch 91, and the weights from epoch 71 were retained as the best epoch. The accuracy was 0.9930 for the training data and 0.9909 for the evaluation data (Fig. 6A), with a loss of 0.0012 for the training data and 0.0016 for the evaluation data (Fig. 6B).

**Figure 6.**
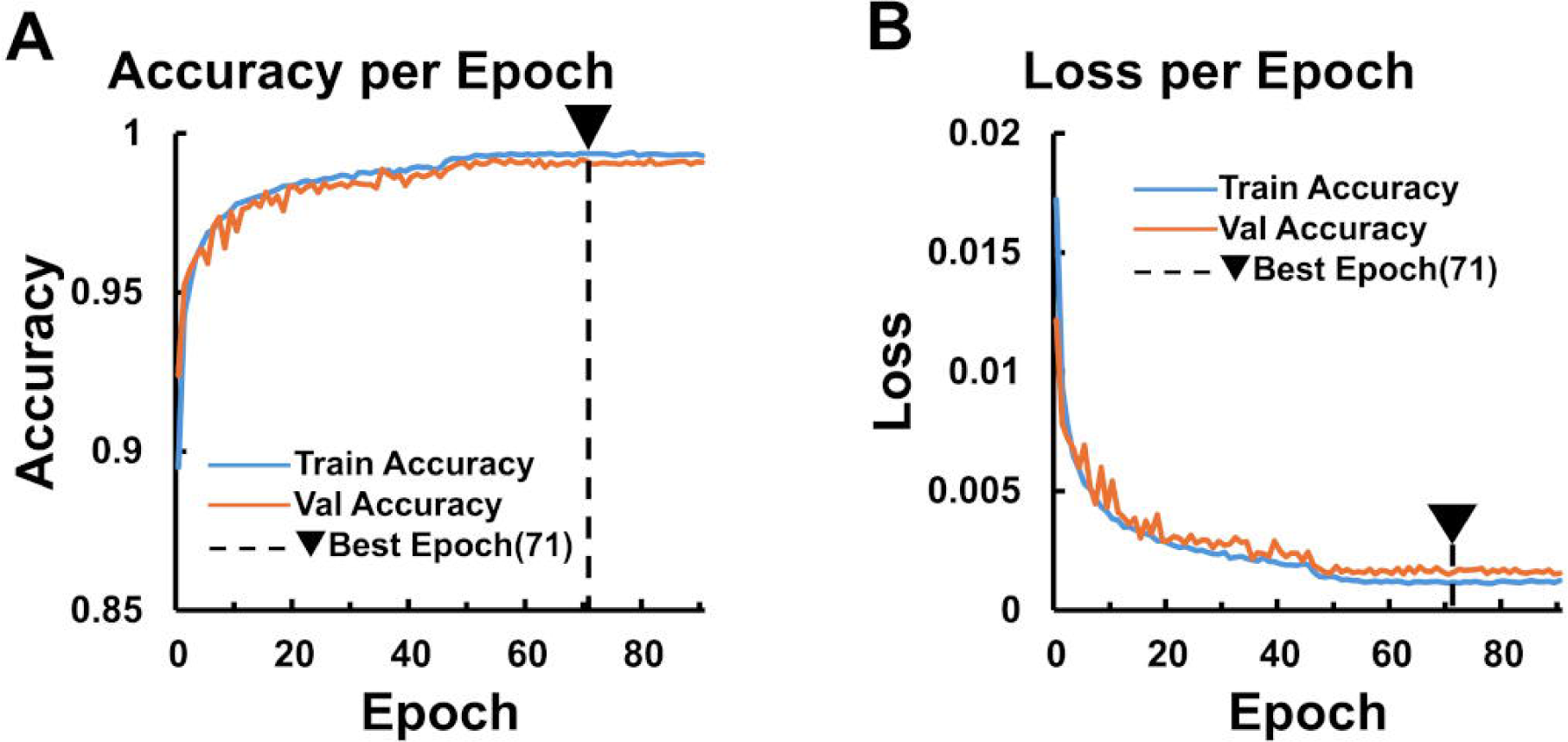
Sex-Determination AI. (A) Accuracy per epoch during training and evaluation. (B) Loss per epoch during training and evaluation. Early stopping was triggered at epoch 91, with the model weights from epoch 71 saved as the best. The results were as follows: for the training data, loss = 0.0012 and accuracy = 0.9930; for the evaluation data, loss = 0.0016 and accuracy = 0.9909.

To evaluate the model, a dataset was predicted to confirm its accuracy, and a confusion matrix was created, as is typical for profile judgment AI. The indices Accuracy, Precision, Recall, and F1 Score were then calculated (Table 4). The model demonstrated an accuracy of 93.87%, indicating that its predictions were overall accurate. For the male evaluation, Precision was 88.99%, meaning that 88.99% of the predicted males were actually male. The Recall rate was 99.80%, indicating that nearly all actual males were correctly predicted as male. Furthermore, the F1 Score, the harmonic mean of Precision and Recall, was 94.09%, reflecting a strong balance between the two metrics. These results suggest that the model achieved a high level of accuracy and robustness in its predictions.

Conversely, in the evaluation of females, the Precision was 99.79%, meaning that 99.79% of those predicted to be female were actually female. The Recall was 88.20%, indicating that 88.20% of actual females were correctly identified as female.

Additionally, the F1 Score, which is the harmonic mean of Precision and Recall, was 93.64%, reflecting a satisfactory outcome. This suggests that, like in males, the model achieves a strong balance between Precision and Recall.

In the sex-determination AI, a video-based evaluation was performed in addition to the previous assessments (Table 5). The Accuracy, Precision, Recall, and F1 Scores all reached 100%, demonstrating excellent performance. This indicates that the application can be used not only for collecting profile images but also for sex determination based solely on profile images.

Grad-CAM++ analysis revealed that the CNN primarily focused on specific regions of the mouse face for sex determination. The activation patterns were similar between male and female mice, with no distinct differences observed. The highlighted regions were concentrated on parts of the face, indicating that the model utilized facial features for classification (Fig. 7).

**Figure 7.**
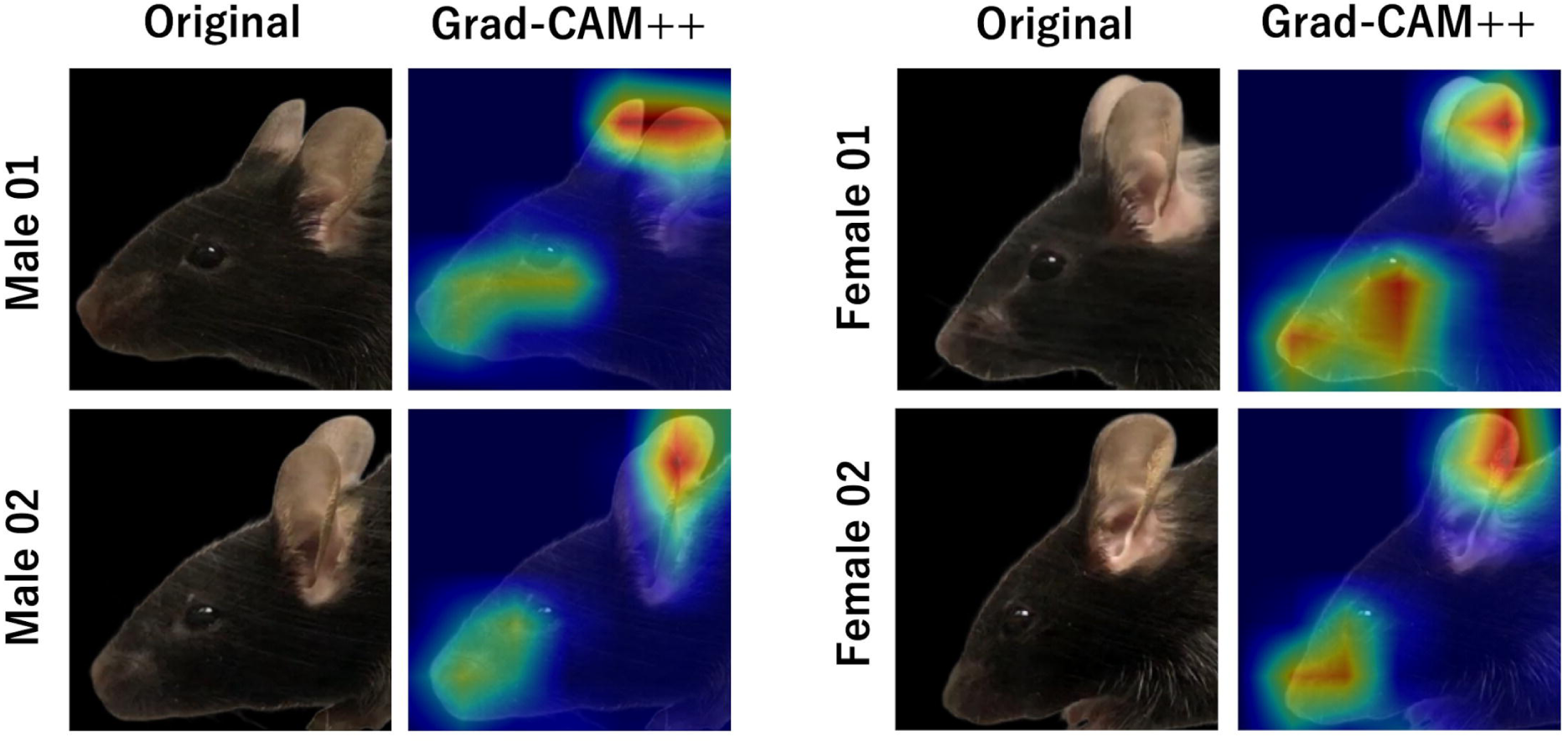
Visualization of the features used by the CNN for sex determination using Grad-CAM++. Grad-CAM++ heatmaps highlight the regions in mouse images that contributed most to the CNN’s decision-making process for distinguishing males and females.

## Discussion

In this study, we developed a deep learning-based application capable of detecting mouse heads with high speed and accuracy, enabling efficient extraction and storage of side-profile images. Two main approaches have been proposed for detecting free-moving rodents: deep learning-based methods and geometric approaches. While geometric approaches are easy to implement and offer rapid processing, they are generally more susceptible to background noise, lighting conditions, and camera artifacts, which can result in inaccurate detection. Deep learning methods, in contrast, have demonstrated higher robustness and accuracy, particularly in complex environments with unstructured backgrounds [28–30].

One of the challenges with existing tools such as Rodent Face Finder is their limited applicability to animals with different coat colors. These tools often rely on brightness contrast and were primarily trained using white rats, making them less reliable for black or brown mice [9,31,32]. To address this limitation, we developed a system trained on animals with varied coat colors, including black, brown, and white mice, as well as *Apodemus speciosus*, thereby extending the tool’s applicability to a broader range of rodent species. Moreover, by automating the final image selection step, we improved the overall efficiency of the data collection pipeline.

We adopted YOLOv8 for head detection due to its well-known balance between speed and accuracy [21]. The model achieved high precision, with mAP50 of 0.995 and mAP50–95 of 0.892, indicating reliable performance across a range of IoU thresholds. Notably, it maintained high accuracy even on test data from mouse strains not included in the training set, such as C3H/He, suggesting that the model has strong generalizability.

To minimize bias related to head orientation, we employed a two-step approach that included a secondary AI for classifying side-profile images. This strategy helped us exclude images with unclear or non-lateral head positions, leading to a more consistent training dataset and potentially more stable inference. The side-profile classification model achieved an accuracy of 91.68%, with a precision of 93.47% and recall of 78.15%. Although some side-profile images were missed, we prioritized high precision to avoid including non-lateral images that could introduce noise into downstream analyses.

However, the use of automated processing on video sequences led to the collection of many visually similar images from consecutive frames, raising concerns about data redundancy. Overrepresentation of near-identical images can bias the model during training and reduce its ability to generalize [33]. To mitigate this, we introduced K-means clustering as a filtering step. By selecting representative images closest to each cluster centroid, we preserved data diversity while avoiding redundancy. Cluster visualizations for K = 3, 8, and 16 revealed that, even at high K values, images appeared similar but differed in posture, orientation, and coat characteristics. This suggests that meaningful diversity was retained within the dataset (Fig. 4E).

To evaluate computational feasibility, we tested the system on both a standard-performance laptop (M1 MacBook Pro) and a high-end RTX-based workstation. In both environments, the application achieved real-time processing speeds, even with video preview enabled. The MacBook Pro achieved 31.2 ms/frame, and the RTX workstation 10.9 ms/frame, both satisfying the threshold of 30 frames per second. Disabling the preview further reduced processing time. In addition, the system supports parallel processing of multiple video files, with practical limits defined by GPU memory capacity—three files for 12 GB and four for 16 GB. These results demonstrate that the application can be deployed flexibly in both field and laboratory settings, including high-throughput experiments.

Beyond head detection, we developed and evaluated a CNN-based AI for sex classification. The model achieved an overall accuracy of 93.87% (Table 4). For males, precision was 88.97% and recall was 99.80%, indicating very few false negatives but a moderate false positive rate. For females, precision reached 99.79%, while recall was lower at 88.20%, meaning some females were missed. F1 scores were well balanced, and track-level evaluation yielded perfect classification accuracy (100%), likely due to the use of consistent side-profile data and the smoothing effect of temporal analysis.

It is notable that the model was able to classify sex with high accuracy in mice, despite their minimal visual differences, especially in inbred strains. This contrasts with species like chimpanzees, where higher inter-individual variation makes visual classification easier [34]. In mice, however, sex-related differences in cranial features—such as the cranial base, vault, and facial bones—have been reported[35]. These differences may subtly influence fur growth patterns and local texture, allowing the CNN to detect minute features not easily perceptible to the human eye.

To investigate this, we conducted Grad-CAM++ analysis [36], which revealed consistent activation in facial areas such as the mouth, whiskers, cheeks, and ears in both males and females. These regions may reflect anatomical structures like the nasal bone, maxilla, zygomatic arch, and premaxilla, suggesting that the model leverages fine-grained textural cues shaped by underlying morphology.

As our model was trained via fine-tuning, it may have focused more on texture-based features than on overall shape [37]. Incorporating end-to-end training in future work may allow the model to learn a more integrated representation of both texture and form, improving robustness.

It is important to note that this study used data from a single mouse strain. Therefore, the model’s performance on other strains remains to be tested. Expanding the training dataset to include multiple strains will be essential to improve generalizability and to explore the tool’s applicability to other rodent species.

Finally, the application’s robustness to background noise and its ability to operate on standard hardware make it a promising tool for ecological research, particularly for non-invasive sex ratio analysis in wild rodents [38]. Further development of automated and generalizable systems will enhance their value in field studies and high-throughput behavioral experiments.

## Acknowledgments

We would like to thank Dr. K. Ohnishi (Department of Pharmacology, Kurume University School of Medicine) for providing the videos of C3H/He and C57BL/6N mice used in this study. A part of this work was conducted at the Institute for Disease Modeling, Kurume University School of Medicine. We gratefully acknowledge the Frontier Science Research Center at the University of Miyazaki for allowing us to use their facilities. Portions of this study were the result of the JSPS KAKENHI (grant number 22K05976).

## References

1. Van Kleef GA, Côté S. The Social Effects of Emotions. Annual Review of Psychology. 2022. doi:10.1146/annurev-psych-020821-010855

2. Chiavaccini L, Gupta A, Chiavaccini G. From facial expressions to algorithms: a narrative review of animal pain recognition technologies. Front Vet Sci. 2024;11. doi:10.3389/fvets.2024.1436795

3. Animal FACS. In: https://animalfacs.com.

4. Waller BM, Peirce K, Caeiro CC, Scheider L, Burrows AM, McCune S, et al. Paedomorphic facial expressions give dogs a selective advantage. PLoS One. 2013;8. doi:10.1371/journal.pone.0082686

5. Mota-Rojas D, Marcet-Rius M, Ogi A, Hernández-Ávalos I, Mariti C, Martínez-Burnes J, et al. Current advances in assessment of dog’s emotions, facial expressions, and their use for clinical recognition of pain. Animals. 2021. doi:10.3390/ani11113334

6. Okano H. Current Status of and Perspectives on the Application of Marmosets in Neurobiology. Annual Review of Neuroscience. 2021. doi:10.1146/annurev-neuro-030520-101844

7. Correia-Caeiro C, Burrows A, Wilson DA, Abdelrahman A, Miyabe-Nishiwaki T. CalliFACS: The common marmoset Facial Action Coding System. PLoS One. 2022;17. doi:10.1371/journal.pone.0266442

8. Langford DJ, Bailey AL, Chanda ML, Clarke SE, Drummond TE, Echols S, et al. Coding of facial expressions of pain in the laboratory mouse. Nat Methods. 2010;7. doi:10.1038/nmeth.1455

9. Sotocinal SG, Sorge RE, Zaloum A, Tuttle AH, Martin LJ, Wieskopf JS, et al. The Rat Grimace Scale: A partially automated method for quantifying pain in the laboratory rat via facial expressions. Mol Pain. 2011;7. doi:10.1186/1744-8069-7-55

10. Keating SCJ, Thomas AA, Flecknell PA, Leach MC. Evaluation of EMLA Cream for Preventing Pain during Tattooing of Rabbits: Changes in Physiological, Behavioural and Facial Expression Responses. PLoS One. 2012;7. doi:10.1371/journal.pone.0044437

11. Steagall P V., Monteiro BP, Marangoni S, Moussa M, Sautié M. Fully automated deep learning models with smartphone applicability for prediction of pain using the Feline Grimace Scale. Sci Rep. 2023;13. doi:10.1038/s41598-023-49031-2

12. Whittaker AL, Liu Y, Barker TH. Methods Used and Application of the Mouse Grimace Scale in Biomedical Research 10 Years on: A Scoping Review. Animals (Basel). 2021;11: 1–27. doi:10.3390/ANI11030673

13. Tuttle AH, Molinaro MJ, Jethwa JF, Sotocinal SG, Prieto JC, Styner MA, et al. A deep neural network to assess spontaneous pain from mouse facial expressions. Mol Pain. 2018;14. doi:10.1177/1744806918763658

14. Le Moëne O, Larsson M. A New Tool for Quantifying Mouse Facial Expressions. eNeuro. 2023;10. doi:10.1523/ENEURO.0349-22.2022

15. Adolphs R, Anderson DJ. The Neuroscience of Emotion: A NEW SYNTHESIS. The Neuroscience of Emotion: A New Synthesis. 2018.

16. Anderson DJ, Adolphs R. A framework for studying emotions across species. Cell. 2014. doi:10.1016/j.cell.2014.03.003

17. Dolensek N, Gehrlach DA, Klein AS, Gogolla N. Facial expressions of emotion states and their neuronal correlates in mice. Science (1979). 2020;368. doi:10.1126/science.aaz9468

18. Yu I, Miyamoto Y, Yoshikawa T, Ohmura Y, Sato M. Facial expression changes in mice during social interaction under head fixation. Proceedings for Annual Meeting of The Japanese Pharmacological Society. 2023;97: 1-B-P-031. doi:10.1254/jpssuppl.97.0_1-b-p-031

19. Tanaka Y, Nakata T, Hibino H, Nishiyama M, Ino D. Classification of multiple emotional states from facial expressions in head-fixed mice using a deep learning-based image analysis. PLoS One. 2023;18. doi:10.1371/journal.pone.0288930

20. Juczewski K, Koussa JA, Kesner AJ, Lee JO, Lovinger DM. Stress and behavioral correlates in the head-fixed method: stress measurements, habituation dynamics, locomotion, and motor-skill learning in mice. Sci Rep. 2020;10. doi:10.1038/s41598-020-69132-6

21. Jocher G, Chaurasia A, Qiu J. YOLO by Ultralytics (Version 8.0.0). Electronics 2020, Vol. 9, Page 1235. 2023.

22. Kuramoto T, Abe S, Ishihata H. Sex Classification of Salmon Using Convolutional Neural Network. Proceedings of the 2020 14th International Conference on Ubiquitous Information Management and Communication, IMCOM 2020. 2020. doi:10.1109/IMCOM48794.2020.9001787

23. Howard A, Sandler M, Chen B, Wang W, Chen LC, Tan M, et al. Searching for mobileNetV3. Proceedings of the IEEE International Conference on Computer Vision. 2019. doi:10.1109/ICCV.2019.00140

24. Lin TY, Goyal P, Girshick R, He K, Dollar P. Focal Loss for Dense Object Detection. IEEE Trans Pattern Anal Mach Intell. 2017;42: 318–327. doi:10.1109/TPAMI.2018.2858826

25. Adrian Rosebrock. Blur detection with OpenCV. https://pyimagesearch.com/2015/09/07/blur-detection-with-opencv/. 7 Sep 2015.

26. Qin X, Zhang Z, Huang C, Dehghan M, Zaiane OR, Jagersand M. U 2-Net: Going Deeper with Nested U-Structure for Salient Object Detection.

27. Daniel Gatis. rembg: Background removal tool. https://github.com/danielgatis/rembg.

28. Kobayashi K, Matsushita S, Shimizu N, Masuko S, Yamamoto M, Murata T. Automated detection of mouse scratching behaviour using convolutional recurrent neural network. Scientific Reports 2021 11:1. 2021;11: 1–10. doi:10.1038/s41598-020-79965-w

29. Kobayashi K, Shimizu N, Matsushita S, Murata T. The assessment of mouse spontaneous locomotor activity using motion picture. J Pharmacol Sci. 2020;143. doi:10.1016/j.jphs.2020.02.003

30. Vidal A, Jha S, Hassler S, Price T, Busso C. Face detection and grimace scale prediction of white furred mice. Machine Learning with Applications. 2022;8. doi:10.1016/j.mlwa.2022.100312

31. Andresen N, Wöllhaf M, Hohlbaum K, Lewejohann L, Hellwich O, Thöne-Reineke C, et al. Towards a fully automated surveillance of well-being status in laboratory mice using deep learning: Starting with facial expression analysis. PLoS One. 2020;15. doi:10.1371/journal.pone.0228059

32. Viola P, Jones M. Rapid object detection using a boosted cascade of simple features. Proceedings of the IEEE Computer Society Conference on Computer Vision and Pattern Recognition. 2001. doi:10.1109/cvpr.2001.990517

33. Mehrabi N, Morstatter F, Saxena N, Lerman K, Galstyan A. A Survey on Bias and Fairness in Machine Learning. ACM Computing Surveys. 2021. doi:10.1145/3457607

34. Schofield D, Nagrani A, Zisserman A, Hayashi M, Matsuzawa T, Biro D, et al. Chimpanzee face recognition from videos in the wild using deep learning. Sci Adv. 2019;5. doi:10.1126/sciadv.aaw0736

35. Percival CJ, Liberton DK, Pardo-Manuel de Villena F, Spritz R, Marcucio R, Hallgrímsson B. Genetics of murine craniofacial morphology: Diallel analysis of the eight founders of the Collaborative Cross. J Anat. 2016;228. doi:10.1111/joa.12382

36. Chattopadhay A, Sarkar A, Howlader P, Balasubramanian VN. Grad-CAM++: Generalized gradient-based visual explanations for deep convolutional networks. Proceedings - 2018 IEEE Winter Conference on Applications of Computer Vision, WACV 2018. 2018. doi:10.1109/WACV.2018.00097

37. Geirhos R, Michaelis C, Wichmann FA, Rubisch P, Bethge M, Brendel W. Imagenet-trained CNNs are biased towards texture; increasing shape bias improves accuracy and robustness. 7th International Conference on Learning Representations, ICLR 2019. 2019.

38. Zemanova MA. Towards more compassionate wildlife research through the 3Rs principles: Moving from invasive to non-invasive methods. Wildlife Biol. 2020;2020. doi:10.2981/wlb.00607

