## Supplemental Table for "AI-Driven System for Large-Scale Automated Collection of Mouse Profile Images"

**Supplement Table 1.** Dataset Pre-Augmentation Multiplication Table.

| Image-augmentations | For side-profile classification AI | | For sex determination AI | |
|---|---|---|---|---|
| | Class “OK” | Class “NG” | Class ”male” | Class “female” |
| Left/right flip | Extended 2-fold | | Extended 2-fold | |
| Random Brightness, Contrast, and Rotation. | Extended 6-fold | Extended 2-fold | Extended 16-fold | |

**Supplemental Table 2.** Dataset Overview for Side-Profile Calssification AI: Image Counts per Class and Totals

| | Class "OK" | Class "NG" | subtotal |
|---|---|---|---|
| C57BL/6N | 3,413 | 6,368 | 9,781 |
| C57BL/6J near-infrared | 54 | 172 | 226 |
| C3H/He | 1,504 | 4,380 | 5,884 |
| ICR | 1,985 | 10,480 | 12,465 |
| Apodemus speciosus | 796 | 853 | 1649 |
| For accuracy evaluation | 1,500 | 4,389 | 5,889 |
| For training | 60,117 | 57,281 | 117,398 |
| For validation | 14,907 | 14,175 | 29,082 |

**Supplemental Table 3.** Dataset Overview for Sex Determination AI: Image Counts per Class and Totals

| | Class "male" | Class "female" | subtotal |
|---|---|---|---|
| Extracted images | 2,694 | 2,697 | 5,391 |
| Cleaned images | 2,675 | 2,670 | 5,345 |
| For accuracy evaluation | 510 | 534 | 1044 |
| For training | 55,462 | 54,720 | 110,182 |
| For validation | 13,818 | 13,632 | 27,450 |

**Supplemantal Table 4.** List of Image Augmentations the Pipeline.

| Image Augmentations Pipeline |
| --- |
| 1. Random zooming in and out. |
| 2. Random left/right flip |
| 3. Random rotation |
| 4. Random translation |
| 5. Random brightness |
| 6. Random contrast |
| 7. Replace the background with gray at a 50% probability, and with random noise at a 50% probability. ※ |

※By replacing the background with gray or random noise at a 50% probability, the model is encouraged to make decisions more based on the object itself [17].

**Supplemental Table 5.** Configuration of MobileNetV3Large Layers for Side-Profile and Sex Determination AI

| Layer | Type | trainable | For side-profile classification AI | For sex determination AI |
|---|---|---|---|---|
| Base model | Headless MobileNetV3Large | Frozen(Up to Conv_1) | Fine-tuned from Conv_1 onward | |
| Global Pooling | Global Average Pooling 2D | Fine-tuned | Included | |
| Convolutional Layer | 1x1 Convolution | Fine-tuned | 1x1x1024 | 1x1x1280 |
| Activation Layer | Activation Function | Fine-tuned | ReLu | Hard-Swish |
| Dropout Layer | Dropout (Rate: 0.2) | Fine-tuned | Included | |
| Logits Layer | Fully Connected Layer | Fine-tuned | Included | |
| Flatten | Flatten Layer | Fine-tuned | Included | |
| Output Layer | SoftMax | Fine-tuned | Included | |
